# A genomic amplification affecting a carboxylesterase gene cluster confers organophosphate resistance in the mosquito *Aedes aegypti*: from genomic characterization to high-throughput field detection

**DOI:** 10.1101/2020.06.08.139741

**Authors:** Julien Cattel, Chloé Haberkorn, Fréderic Laporte, Thierry Gaude, Tristan Cumer, Julien Renaud, Ian W. Sutherland, Jeffrey C. Hertz, Jean-Marc Bonneville, Victor Arnaud, Camille Noûs, Bénédicte Fustec, Sébastien Boyer, Sébastien Marcombe, Jean-Philippe David

## Abstract

By altering gene expression and creating paralogs, genomic amplifications represent a key component of short-term adaptive processes. In insects, the use of insecticides can select gene amplifications causing an increased expression of detoxification enzymes, supporting the usefulness of these DNA markers for monitoring the dynamics of resistance alleles in the field. In this context, the present study aims to characterise a genomic amplification event associated with resistance to organophosphate insecticides in the mosquito *Aedes aegypti* and to develop a molecular assay to monitor the associated resistance alleles in the field. An experimental evolution experiment using a composite population from Laos supported the association between the over-transcription of multiple contiguous carboxylesterase genes on chromosome 2 and resistance to multiple organophosphate insecticides. Combining whole genome sequencing and qPCR on specific genes confirmed the presence of a ~100 Kb amplification spanning at least five carboxylesterase genes at this locus with the co-existence of multiple structural duplication haplotypes. Field data confirmed their circulation in South-East Asia and revealed high copy number polymorphism among and within populations suggesting a trade-off between this resistance mechanism and associated fitness costs. A dual-colour multiplex TaqMan assay allowing the rapid detection and copy number quantification of this amplification event in *Ae. aegypti* was developed and validated on field populations. The routine use of this novel assay will improve the tracking of resistance alleles in this major arbovirus vector.

## Introduction

The amplification of genomic regions overlapping genes can affect fitness by creating paralogs that can diverge to generate novel functions or by altering gene expression levels (Iskow et al., 2012). Often called genomic amplification when multiple copies are present, this mechanism has been shown to be a major driver of short-term adaptation (Kondrashov, 2012). Indeed, amplification events are frequent in natural populations (Campbell & Eichler, 2013) and can be subjected to positive selection (Chatonnet et al., 2017; Cooper et al., 2007; Zhang, 2003).

In insects, genomic amplifications have been shown to play a key role in the evolution of insecticide resistance by three distinct mechanisms. First, the duplication of genes encoding insecticide targets can allow resistant individuals to reduce the fitness costs associated with target-site mutations by allowing the co-existence of the susceptible and the resistant alleles (Assogba et al., 2015; Labbé et al., 2007). Second, the neofunctionalization of new gene copies can lead to novel adaptive functions (Zimmer et al., 2018). Third, genomic amplifications affecting genes encoding detoxification enzymes leading to their over-expression can confer the insect a higher capacity to degrade or sequester insecticides (Bass & Field, 2011). This latter mechanism has been reported in all three mosquito genera of high medical importance, *Aedes*, *Anopheles* and *Culex*, affecting various detoxification enzyme families including cytochrome P450 monooxygenases (*P450*s) and carboxy/choline esterases (*CCE*s) (Cattel et al., 2019; Lucas et al., 2019; Weetman et al., 2018).

Genomic amplifications affecting *CCE*s have even been described as the most common route of enzyme over-production in mosquitoes (Bass & Field, 2011). A classic example comes from the house mosquito *Culex pipiens* in which amplified carboxylesterases genes (in conjunction with the *ace 1* G119S target-site mutation) have spread across the globe, providing high resistance to organophosphate insecticides (Raymond et al., 2001). In *Aedes* mosquitoes, the low chance of occurrence of the G119S *ace-1* mutation because of strong genetic constraints (Weill et al., 2004) suggest that *CCE* amplifications play a central role in organophosphate resistance and are thus of high interest for resistance monitoring.

In the tiger mosquito *Aedes albopictus*, the over-expression of two *CCE* genes (*CCEAE3A* and *CCEAE6A*) through gene amplification was associated with resistance to the organophosphate insecticide temephos (Grigoraki et al., 2017). In the yellow fever mosquito *Aedes aegypti*, AAEL023844 (formerly *CCEAE3A*) and other *CCE* genes belonging to the same genomic cluster were also found overexpressed through gene amplification in temephos-resistant populations (Faucon et al., 2015, 2017; Poupardin et al., 2014). Further functional studies confirmed that *CCEAE3A* is able to sequester and metabolize the active form of temephos in both *Ae. aegypti* and *Ae. albopictus* (Grigoraki et al., 2016). Although the genomic structure and polymorphism of this *CCE* amplification was studied in *Ae. albopictus* (Grigoraki et al., 2017) such work has not been conducted in *Ae. aegypti*. In addition, the role of *CCEAE3A* amplification in resistance to other insecticides remains unclear. Finally, no high-throughput assay has yet been developed to track this resistance mechanism in natural populations although such a tool would significantly ease resistance monitoring and management.

In this context, we combined an experimental evolution experiment with RNA-seq and whole genome sequencing to confirm the association between this genomic amplification, the overexpression of *CCE* genes and resistance to the organophosphate insecticide malathion in *Ae. aegypti*. Bioassay data also support the importance of the *CCE* amplification in resistance to other organophosphates insecticides. Comparing gene Copy Number Variations (CNV) between the different genes of this genomic cluster suggested the presence of at least two distinct haplotypes occurring in South-East Asia (SEA), both associated with resistance. Investigating their spatial dynamics in natural populations confirmed their co-occurrence in the field with a high copy number polymorphism within and among populations. Based on these results, we developed a novel high-throughput multiplex TaqMan assay allowing the quantitative detection of this *CCE* amplification in hundreds of individual mosquitoes within a few hours. By reducing the human power and infrastructure needs associated with bioassays, this molecular assay will facilitate the tracking of organophosphate resistance alleles in natural *Ae. aegypti* populations.

## Material and methods

### Field sampling and mosquito lines

*Aedes aegypti* larvae and pupae were collected in households and temples of eleven villages belonging to five provinces of Laos in 2014 (Table S1). Previous work confirmed the circulation of organophosphate and pyrethroid resistance alleles in these populations together with the presence of amplification affecting AAEL023844 (formerly *CCEAE3A*) (Marcombe et al., 2019). These populations were maintained under controlled conditions (27 ± 2 °C and 80 ± 10% relative humidity) at the Institut Pasteur of Laos for 5 generations and used for experimental evolution (see below). A second round of larvae collection was conducted in 2017 for studying the spatial dynamics of *CCE* genomic amplifications in SEA. Fourteen different populations were sampled in Laos, Thailand and Cambodia (see details in Table S1) and adults were stored individually at −20°C in silica gel until molecular analyses.

### Experimental selection

A Laos composite population was created by pooling 50 virgin males and 50 virgin females of each population into a single cage (Table S1). This population was then maintained for 2 generations without insecticide selection to allow genetic mixing before initiating insecticide selection. The Laos composite population was then split in 2 lines (N > 1000 in each line): one line being maintained without insecticide selection (NS line) while the second line (Mala line) was artificially selected with malathion at the adult stage for 4 consecutive generations (from G1 to G5). For this, batches of thirty-three-day-old non-blood-fed adult mosquitoes (~1000 individuals of mixed sex) were exposed at each generation to filters papers impregnated with malathion using WHO test tubes. A constant dose of 5% malathion coupled with an exposure time of 10 min (leading to ~90% mortality at G1) were used through the whole selection process. Surviving adults (mainly females) were collected 48h after insecticide exposure, blood fed on mice and allowed to lay eggs to generate the next generation. Three-day-old non-blood-fed adult females (not exposed to insecticide) were sampled after four generations and used for bioassays and molecular work. Mosquitoes were identified as follows. G1: initial composite population, G5-NS: line maintained without insecticide pressure for four generations, G5-Mala: line maintained under malathion selection for four generations. Sampled mosquitoes were stored at −20°C until molecular analyses.

### Bioassays

Bioassays were used to monitor the dynamics of malathion resistance during the selection process. Four replicates of 20 three-day-old non-blood-fed females not previously exposed to insecticide and reared in same insectary conditions were sampled at each generation and exposed to the insecticide as described above using the same dose and exposure time as for artificial selection. Mortality was recorded 48h after exposure. Cross resistance to other insecticides was investigated in G5 individuals (G5-Mala and G5-NS) not previously exposed to insecticide. Calibrated individuals were exposed to three distinct insecticides: the organophosphates fenitrothion and temephos, and the pyrethroid deltamethrin. For the adulticides fenitrothion and deltamethrin, bioassays were performed on eight replicates of fifteen three-day-old non-blood-fed females with the following doses and exposure times: fenithrotion 1% for 30 min, deltamethrin 0.05% for 20 min. Mortality rates were recorded 48h after exposure. For the larvicide temephos, bioassays were performed on eight replicates of twenty calibrated third instar larvae exposed to 0.08 mg/μL temephos for 24h in 200 ml tap water and mortality was recorded at the end of exposure.

### RNA sequencing

RNA-sequencing was performed to compare gene expression levels between the NS line, the Mala line and the fully susceptible reference line Bora-Bora. This experiment was performed on unexposed G6 individuals (progeny of the last generation of selection) in order to avoid gene induction/repression effects that can be caused by insecticide exposure. For each line, four RNA-seq libraries were prepared from distinct batches of 25 calibrated three-day-old non-blood-fed females not exposed to insecticide. Total RNA was extracted using Trizol^®^ (Thermo Fisher Scientific) following manufacturer’s instructions. RNA samples were then treated with the RNase-free DNase set (Qiagen) to remove gDNA contaminants and QC checked using Qubit (Thermo Fisher Scientific) and bioanalyzer (Agilent). RNA-seq libraries were prepared from 500 ng total RNA using the NEBNext^®^ Ultra™ II directional RNA library Prep Kit for Illumina (New England Biolabs) following manufacturer’s instructions. Briefly, mRNAs were captured using oligodT magnetic beads and fragmented before being reverse transcribed using random primers. Double stranded cDNAs were synthesized end-repaired and adaptors were incorporated at both ends. Libraries were then amplified by PCR for 10 cycles and purified before QC check using Qubit fluorimeter and Bioanalyzer. Libraries were then sequenced in multiplex as single 75 bp reads using a NextSeq500 sequencer (Illumina). After unplexing and removing adaptors, sequenced reads from each library were loaded into Strand NGS V3.2 (Strand Life Science) and mapped to the latest *Ae. aegypti* genome assembly (Aaeg L5) using the following parameters: min identity = 90%, max gaps = 5%, min aligned read length = 25, ignore reads with >5 matches, 3’end read trimming if quality <20, Kmer size = 11, mismatch penalty = 4, gap opening penalty = 6, gap extension penalty = 1. Mapped reads were then filtered based on their quality and alignment score as follows: mean read quality > 25, max N allowed per read = 5, mapping quality ≥120, no multiple match allowed, read length ≥ 35. Quantification of transcription levels was performed on the 14626 protein-coding genes using the DESeq method with 1000 iterations (Anders & Huber, 2010). Only the 11825 genes showing a normalized expression level ≥ 0.5 (~0.05 RPKM) in all replicates for all lines were retained for further analysis. Differential gene transcription levels between each line across all replicates were then computed using a one-way ANOVA followed by a Tukey post-hoc test and P values were corrected using the Benjamini and Hockberg multiple testing correction (Benjamini & Hochberg, 1995). Genes showing a fold change (FC) ≥ 3 (in either direction) and a corrected P value ≤ 0.001 in the G6-Mala line versus both susceptible lines (G6-NS and Bora-Bora) were considered as differentially transcribed in association with insecticide resistance.

### Whole genome sequencing

The occurrence of a genomic amplification affecting the *CCE* cluster on chromosome 2 at ~174 Mb was investigated by sequencing the whole genome of the Nakh-R population from Thailand as compared to the fully susceptible line Bora-Bora. This population was used for the genomic characterization of this CCE amplification because *i*) this population was known resistant to carry organophosphate resistance alleles (Faucon et al., 2015), *ii*) it showed an over-expression of this CCE gene cluster likely associated with a genomic amplification (Faucon et al., 2015, 2017) *iii*) it was collected from the field and therefore can be used to control for genetic drift effects potentially occurring in the laboratory selected line (G5-Mala). For each population, genomic DNA was extracted from 2 batches of 50 adult females and gDNA extracts were then pooled in equal proportion into a single sequencing library as described in Faucon et al., (2015). Whole genome sequencing was performed from 200 ng gDNA. Sequencing libraries were prepared according to the TruSeq DNA Nano Reference guide for Illumina Paired-end Indexed sequencing (version oct 2017) with a mean insert size of 550 bp. Sequencing was performed on a NextSeq 550 (Illumina) as 150 bp paired-reads. After unplexing and adaptor removal, reads were mapped to the latest *Ae. aegypti* genome assembly (Aaeg L5) using BWA-MEM with default parameters (version 0.7.12). Sequenced reads were then sorted using samtools sort (version 1.2), annotated using Picard FixMateInformation (version 1.137) and PCR duplicates were identified using Picard MarkDuplicates (version 1.137). Normalized coverage profiles between the resistant and the susceptible populations were then compared using non-duplicated reads with a mapping quality score above 60.

### Quantification of Copy Number Variations

Among the six genes located within the genomic amplification detected on chromosome 2, three genes clearly annotated as *CCE* and distributed throughout the cluster were studied: AAEL019678, AAEL023844 (formerly *CCEAE3A*) and AAEL005113 (*CCEAE1A*). For each gene, specific primer pairs were designed using NCBI primer Blast (Table S2). In order to quantify CNV in natural populations and in individuals, genomic DNA was extracted either from seven pools of five adult females (mean CNV comparison between lines) or from single adult females (estimation of amplification prevalence) using the cetyltrimethylammonium bromure (CTAB) method (Collins et al., 1987) and diluted to 0.5 ng/μL prior to amplification. Pooled samples were amplified in duplicates while individual mosquito samples were amplified only once. Quantitative PCR reactions consisted of 3 μL gDNA template, 3.6 μL nuclease free water, 0.45 μL of each primer (10μM), and 7.5 μL of iQ SYBR Green Supermix (Bio-Rad). PCR amplification were performed on a CFX qPCR system (Bio-Rad) with cycles as follows: 95°C 3 min followed by 40 cycles of 95°C 15 secs and 30 secs for hybridization. A dilution scale made from a pool of all gDNA samples was used for assessing PCR efficiency. Data were analyzed using the ΔΔCt method (Pfaffl, 2001) taking into account PCR efficiency. Two control genes (the *P450* AAEL007808 and the chloride channel protein AAEL005950) shown to be present as single copies in multiple *Ae. aegypti* strains and populations (Faucon et al., 2015) were used for normalization. For each gene, CNV were expressed as mean relative gDNA quantity as compared to the fully susceptible line Bora-Bora. For assessing genomic amplification frequencies, all individuals showing a CNV ≥ 2.5-fold as compared to the Bora-Bora line were considered positive. This threshold was chosen in order to avoid false positives as a consequence of qPCR technical variations (< 2-fold in negative control samples). Structural duplication haplotypes were assigned based on the detection of CNV for all three CCE genes (haplotype A) or only for AAEL019678, and AAEL023844 (haplotype B). Individual CNV levels obtained for the *CCE* gene AAEL023844 by qPCR were cross-validated by digital droplet PCR (ddPCR). Briefly, each sample was partitioned into ~20,000 nanoliter-sized droplets using the QX 200 droplet generator (Bio-Rad) by mixing synthetic oil with 20 μL PCR mix containing 2X ddPCR Evagreen supermix (Bio-Rad), 0.9 mM of each primer and 5 μL of template gDNA at 0.5 ng/μL. Emulsified reaction mixtures were then amplified with a C1000 thermal cycler (Bio-Rad) for 40 cycles. After amplification, the number of positive and negative droplets were quantified for each sample using the QX 200 droplet reader (Bio-Rad) and the positive/negative ratio was used to estimate the initial DNA assuming a Poisson distribution. A similar procedure was applied to the control gene AAEL007808 present as a single copy. After normalizing for initial gDNA quantity, CNV were expressed as relative gDNA quantity as compared to the fully susceptible line Bora-Bora.

### CNV quantification using TaqMan multiplex assay

A TaqMan multiplex assay allowing the concomitant quantification of the *CCE* gene AAEL023844 (present in both duplication haplotypes) and the control gene AAEL007808 from single mosquitoes within the same qPCR reaction was developed. For each gene, primers and probes were designed using Primer3web v 4.1.0 (Rozen & Skaletsky, 2000) with the AaegL5 assembly as reference genome for assessing specificity. For each gene, exonic regions were targeted in order to limit amplification variations associated with natural polymorphism (Table S2). The assay was then tested on all individuals detected as positive by qPCR representing 27 individuals belonging to seven populations from three countries. Each reaction mixture contained 12.5 μL of qPCR probe Master Mix (Bio-Rad), 2.25 μL of each primer (10μM), 0.625 μL of each probe (10 μM), 1.25 μL of nuclease free water and 1 μL of template DNA (0.5 ng/μL). PCR amplifications were performed on a CFX qPCR system (Bio-Rad) with cycles set as follows: 95°C for 10 min followed by 40 cycles of 95°C for 10 secs and 60°C for 45 secs followed by FAM and HEX levels reading (see Supplementary File 1 for a user guideline on this TaqMan assay).

### Statistical analysis

All statistical analyses were performed with R v3.6.2 (R Core Team, 2013), using the package lme4 for all mixed models (Bates et al., 2015). Mortality data were statistically compared across conditions by using a Generalized Linear Model (GLM) with mixed effects (binomial family) in which the replicates were included as a random factor. For comparing mean CNV obtained from pools of mosquitoes, normalized gDNA levels obtained for each gene were Log2 transformed and compared across conditions using a GLM with mixed-effects in which the replicates were included as a random factor. For comparing CNV obtained from individual mosquitoes, normalized gDNA levels were Log2 transformed and compared between genes using a one-way ANOVA. A Pearson’s product moment correlation coefficient test was used to compare normalized gDNA quantities obtained from qPCR and ddPCR.

## Results

### Dynamics of organophosphate resistance during experimental selection

Maintaining the Laos composite population under selection with malathion resulted in the rapid rise of resistance (Figure 1).

**Figure 1.**
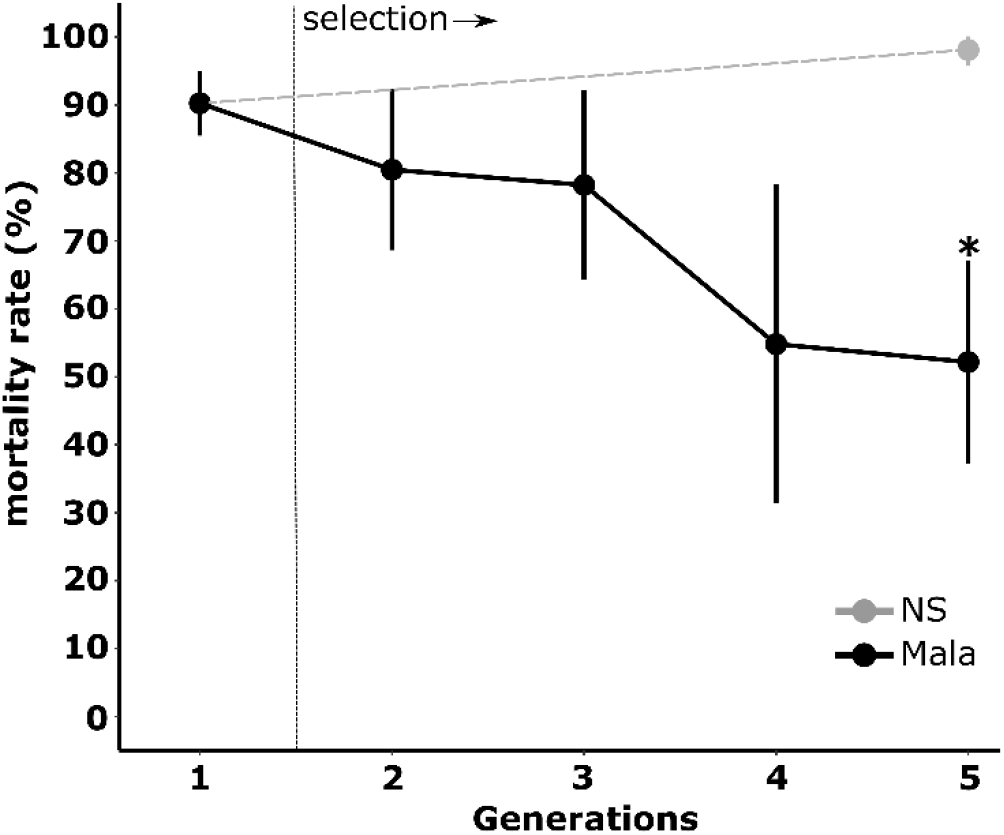
Dynamics of malathion resistance during the selection process. Black: Laos composite population selected with malathion; Grey: Laos composite population maintained without selection. Stars indicate a significant mortality difference as compared to the initial population (N=4, GLM mixed effects binomial family, *p<0.05).

Mortality to malathion dropped gradually from 90.3% in G1 to 52.2 % after four generations of selection (GLMER test: z=2.058, P<0.05 for G5-Mala vs G1). Conversely, we observed a slight (not significant) increase of mortality to 99.1% after four generations without selection (GLMER test: z=−0.433, P=0.665). Bioassays performed with different insecticides revealed that selection with malathion for four generations also select resistance to other organophosphate insecticides (Figure 2).

**Figure 2.**
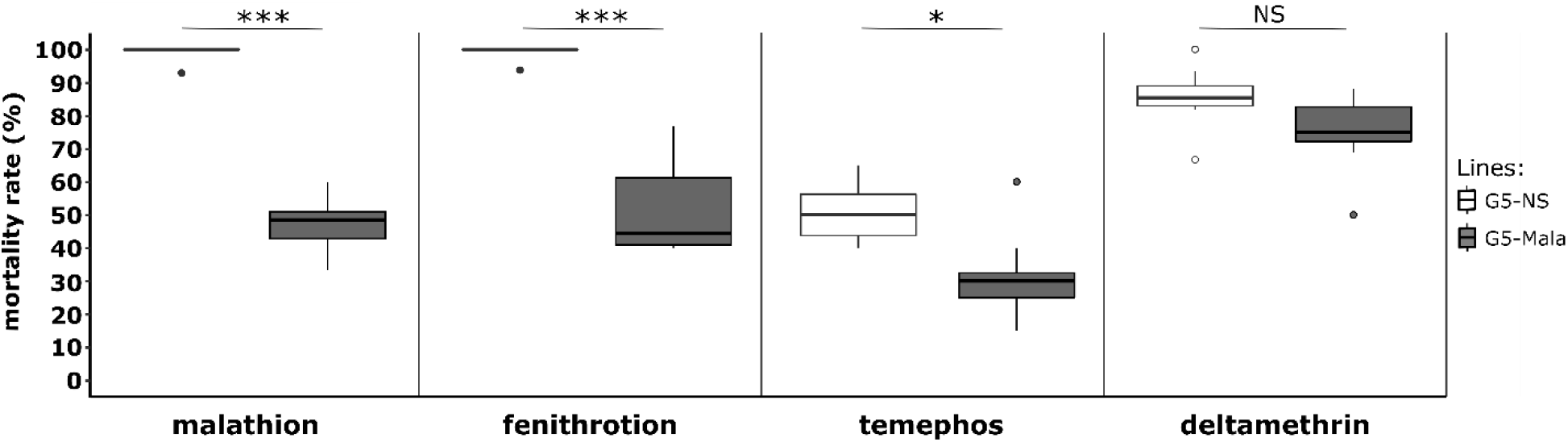
Cross resistance of the Malathion-resistant line to other insecticides. G5-NS: Composite population maintained without selection. G5-Mala: composite population selected with malathion. For each insecticide, stars indicate a significant mortality difference between G5-NS and G5-Mala individuals (N=8, GLM mixed effects binomial family, *p<0,05, ***p<0,001, NS: non-significant).

As compared to the G5-NS line, the G5-Mala line showed a significant increased resistance to the organophosphates fenitrothion at the adult stage (GLMER: z=−3.455, P<0.005) and temephos at the larval stage (GLMER: z=−2.194, P<0.05). Conversely, no significant increased resistance to the pyrethroid deltamethrin at the adult stage was observed suggesting that malathion did not select for pyrethroid resistance alleles.

### Genes associated with malathion resistance

RNA-seq analysis identified 84 and 124 genes over- and under-transcribed, respectively, in G6-Mala adult females (adjusted P value ≤ 0.001 and FC ≥ 3-fold) as compared to the susceptible lines G6-NS and Bora-Bora (Table S3). Among them, 24 genes encoded proteins potentially associated with known insecticide resistance mechanisms (target-site resistance, cuticle alteration, detoxification and sequestration or altered transport) including 14 cuticle genes and 10 detoxification genes. Only seven candidate genes were over-transcribed in the G5-Mala line, all being associated with detoxification (Figure 3A). This included a microsomal glutathione S-transferase on chromosome 1 (AAEL006818, 13-fold *versus* G6-NS), the cytochrome P450 *CYP6N17* on chromosome 2 (AAEL010158, 8-fold *versus* G6-NS) and five contiguous *CCE* genes at ~174 Mb on chromosome 2 (AAEL015304, AAEL019679, AAEL019678, AAEL005123 and AAEL023844 (formerly *CCEAE3A*, up to 10-fold *versus* G6-NS). A closer look at this genomic region revealed the presence of an additional *CCE* gene on the 5’ side of the cluster (AAEL005113 *CCEAE1A*) which was not significantly over-transcribed in the resistant line. Although the *GST* AAEL006818 and the P450 *CYP6N17* were significantly over-transcribed in the G6-Mala line, these two genes were also found over-transcribed in two other Laos lines selected with unrelated insecticides (data not shown), and may thus not be specially associated with malathion resistance.

**Figure 3.**
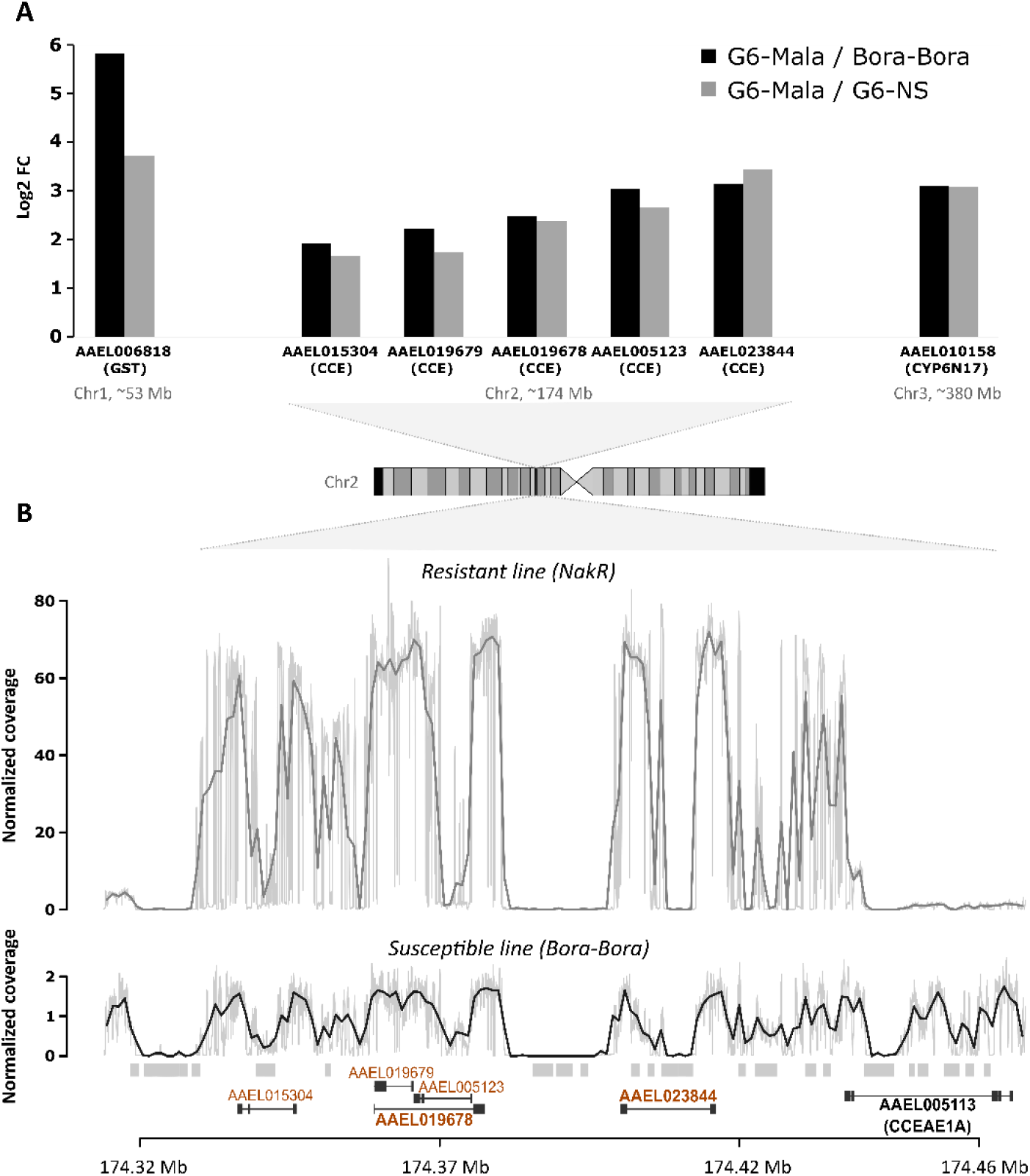
Genomic amplification associated with carboxylesterase overexpression. A: Detoxification genes over-transcribed in the malathion-resistant line. Transcription levels were quantified by RNA-seq using 4 biological replicates per line. Detoxification genes showing a 3-fold over-transcription and a corrected P value <0.001 in the Malathion-resistant line G6-Mala versus both the unselected line NS (grey) and the fully susceptible line Bora-Bora (black) are shown. The genomic location of each gene is indicated. B: Comparison of normalized read coverage profiles at the carboxylesterase locus between a Thai resistant line (Nakh-R) (grey) and the fully susceptible line Bora-Bora (black). Coverage profiles were obtained by divided the raw coverage by the average coverage found on the chromosome 1 and from whole genome DNA-seq performed on pools of 100 individuals. The genomic location of carboxylesterase genes is indicated according to Aaeg L5.1 annotation. Genes found overexpressed by RNA-seq are shown in orange. Genes targeted by qPCR are shown in bold. Repeated elements associated with coverage gaps are indicated as grey boxes.

Cross-comparing RNA-seq data with normalized coverage profiles obtained from whole genome sequencing of the Thai resistant population Nakh-R known to over-express these CCE genes (Faucon et al., 2015, 2017) revealed a ~50-fold increased coverage in this genomic region as compared to the susceptible line (Figure 3B). In addition, this region included multiple low-coverage sections associated with the presence of repeated elements (mostly due to unresolved read assembly). The pattern of genomic amplification observed in the Thai resistant population Nakh-R was in agreement with the expression pattern observed in the Laos resistant line G6-Mala with the same five first CCE genes being amplified, but not *CCEAE1A*.

The occurrence of this genomic amplification in Laos was then confirmed by qPCR (Figure 4A). Before selection (G1), a slight non-significant elevation of gene copy number was observed for three *CCE* genes belonging to this genomic cluster as compared to the fully susceptible line Bora-Bora with important variations suggesting a high inter-individual heterogeneity in the initial line. Although not significant, gene copy number were even lower after four generations without selection (G5-NS) with less variations observed. Conversely, four generations of selection with malathion lead to a strong increase in gene copy number for the *CCE* genes AAEL019678 and AAE023844 in G5-Mala individuals (up to 32-fold). A lower increase (~8-fold) associated with a higher variance was observed for the gene *CCEAE1A*, suggesting that not all G5-Mala individuals carry multiple copies of this gene.

**Figure 4.**
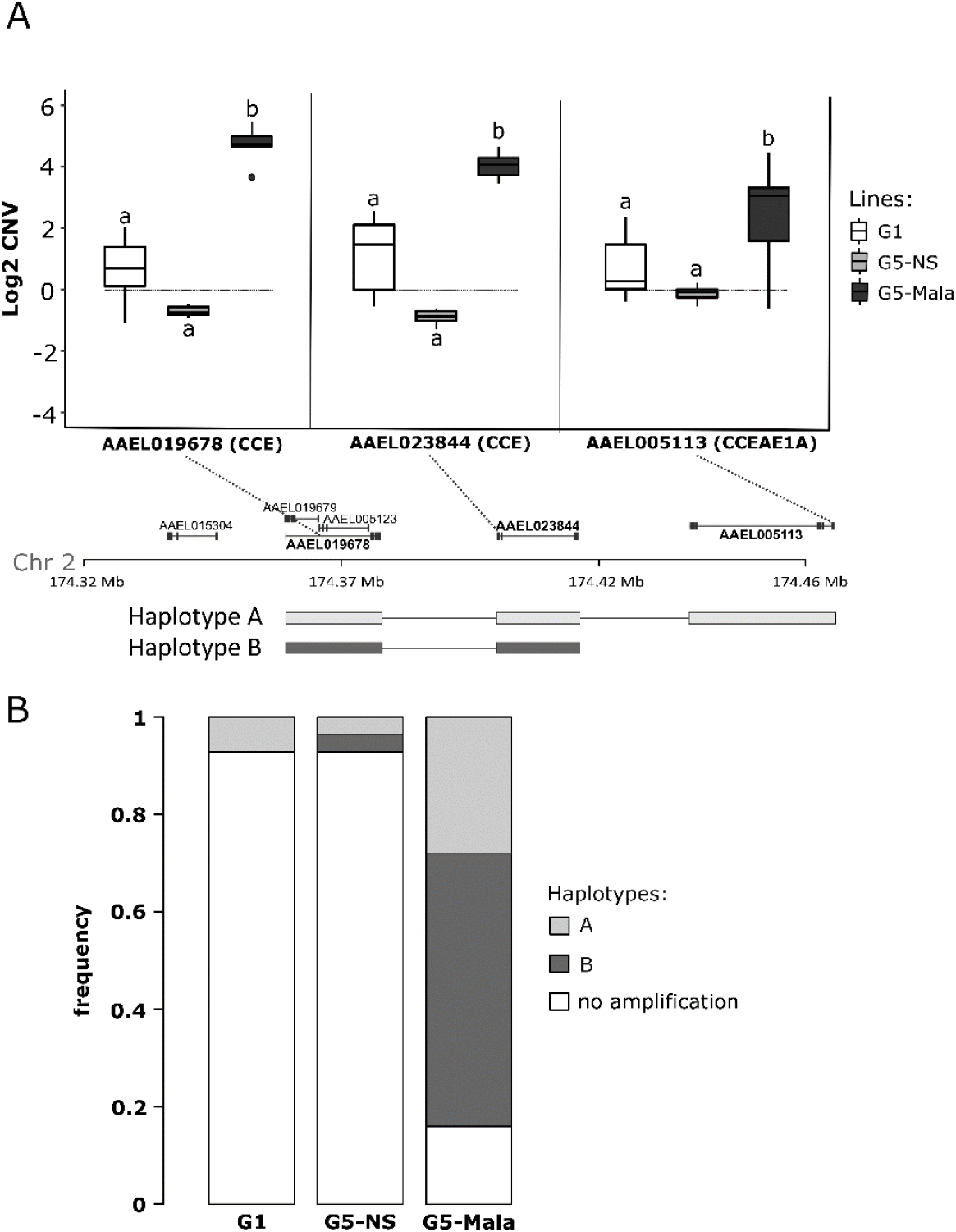
Duplication haplotypes at the carboxylesterase locus. A: CNV of three selected CCE genes located in different positions of the carboxylesterase locus. For each gene, the position of the qPCR amplification product is indicated (dashed lines). Mean CNV were estimated by qPCR on pools of mosquitoes and are expressed as gDNA quantity relative to the fully susceptible line Bora-Bora (horizontal grey line). G1: unselected line at generation 1, G5-NS: unselected line at generation 5, G5-Mala: malathion-resistant line after 4 generations of selection. Distinct letters indicate significant mean CNV variations between populations (GLM mixed effects, N=7, p≤ 0.05). The two structural duplication haplotypes deduced from CNV data are represented. B: Frequencies of each haplotype in the different lines. Haplotypes frequencies were deduced from individual CNV data obtained by qPCR from 28 mosquitoes per line for the three genes AAEL019678, AAEL023844 (formerly *CCEAE3A*) and AAEL005113 (*CCEAE1A*).

Quantification of *CCE* genes copy number in individual mosquitoes by qPCR confirmed the presence of at least two distinct structural duplication haplotypes in Laos with haplotype A including the three *CCE* genes and haplotype B not including *CCEAE1A* (Figure 4B). While the prevalence of these two *CCE* haplotypes was low in the initial composite population (7% for the haplotype A and 0% for the haplotype B) and in the G5-NS (<4% for each haplotype) their cumulated frequency reached 84% in G5-Mala individuals with haplotype B being more frequent (at least 67% of duplicated haplotypes) than haplotype A (33% of duplicated haplotypes).

### Prevalence and copy number polymorphism in SEA

The spatial dynamics of this genomic amplification event was investigated in field populations from Laos, Cambodia and Thailand. A total of 302 mosquitoes belonging to 14 field populations were genotyped for the presence of *CCE* amplifications using qPCR. Seven populations distributed across the three countries were found positive with at least one individual carrying the duplication haplotype A or B (Figure 5). The prevalence of *CCE* amplifications was low in most studied populations, high in two populations from Cambodia (26% and 29%) and very high in the Nakh-R Thai population known to be resistant to organophosphates (79 %) (Faucon et al., 2015). Although both haplotypes were detected through the study area, all positive individuals from populations showing a high prevalence only carried haplotype B.

**Figure 5.**
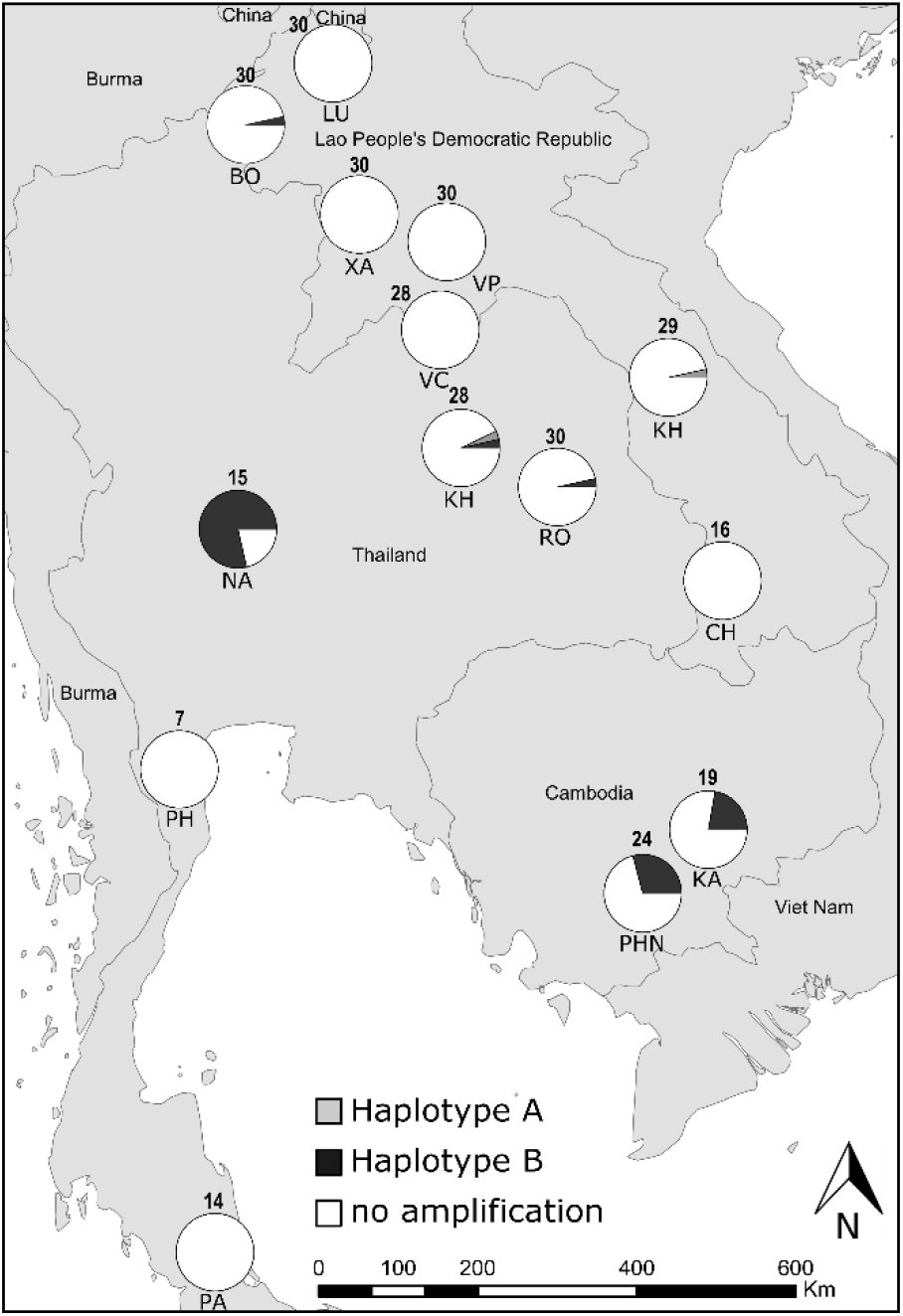
Prevalence of the carboxylesterase gene amplification in South-East Asia. For each population, the frequency of each haplotype is shown. The number of individuals genotyped is shown above each pie chart. Haplotypes frequencies were deduced from CNV data obtained by qPCR on individual mosquitoes for the three genes AAEL019678, AAEL023844 and AAEL005113 (*CCEAE1A*). Only individuals showing a CNV ≥ 2.5-fold (as compared to the susceptible line Bora-Bora) for any gene were considered positive. The description of the studied populations is presented in Table S1.

Cross comparing CNV data obtained for the *CCE* gene AAEL023844 between standard Sybrgreen qPCR and digital droplet PCR (ddPCR) indicated a good correlation between the two techniques (r=0.85, P<0.001, Figure S1), suggesting that despite the technical variations inherent to qPCR on single mosquitoes this approach provides a relatively good estimation of gene copy numbers. Comparing the number of copies of each *CCE* gene between all positive individuals revealed an important copy number polymorphism in the SEA with estimated copy numbers ranging from 3 to ~80 copies (Figure 6). No significant differences were observed between field populations (P=0.578 for AAEL019678; P=0.721 for CCEAE3A) and the different lines (G1, G5-NS, G5-Mala) (P=0.827 for AAEL019678; P=0.845 for CCEAE3A), suggesting that insecticide selection rather select for positive individuals than for individuals carrying a higher number of copies. Overall, the mean copy number observed for the *CCE* gene AAEL023844 (present in both haplotypes) was significantly higher than for the two other *CCE* genes AAEL019678 and *CCEAE1A* (P<0.001 and P<0.01 respectively) possibly reflecting additional structural haplotypes affecting this gene.

**Figure 6.**
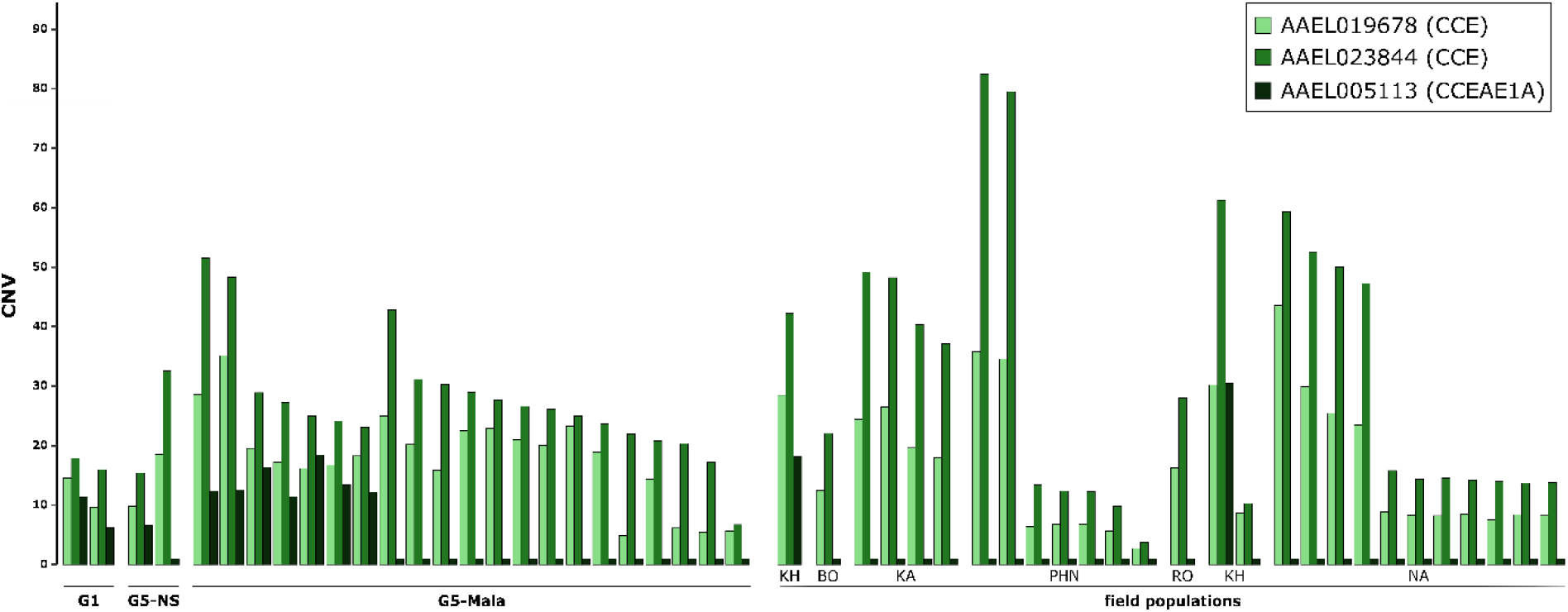
Genes copy number variations in experimental lines and field populations. For each gene, CNV were estimated by qPCR on individual mosquitoes and are expressed as gDNA quantity relative to the fully susceptible line Bora-Bora. Only positive individuals showing a CNV ≥ 2.5-fold for any gene are shown. Names of laboratory lines and field populations are indicated.

### A novel TaqMan assay to track *CCE* gene amplification in *Ae. Aegypti*

A multiplex TaqMan qPCR assay allowing the concomitant amplification of the *CCE* gene AAEL023844 (included in both haplotypes) and a control gene within a single reaction was developed and tested against 27 positive individuals and 7 negative individuals belonging to all populations from which *CCE* amplification were detected (Figure 7). This assay showed a good specificity and sensitivity for detecting *CCE* amplifications in *Ae. aegypti*. All samples identified as positive by qPCR (i.e. showing a CNV higher than 2.5-fold) were also identified as positive using the TaqMan assay (no false negatives) and no false positives was observed. A good amplification specificity was observed for both the *CCE* gene AAEL023844 and the control gene. A similar PCR efficiency of ~ 95% was observed for both the *CCE* gene AAEL023844 and the control gene leading to a Cq of ~30 cycles for both genes in absence of amplification with 0.5 ng/μL template gDNA (see Supplementary File 1 for a user guideline on this TaqMan assay).

**Figure 7.**
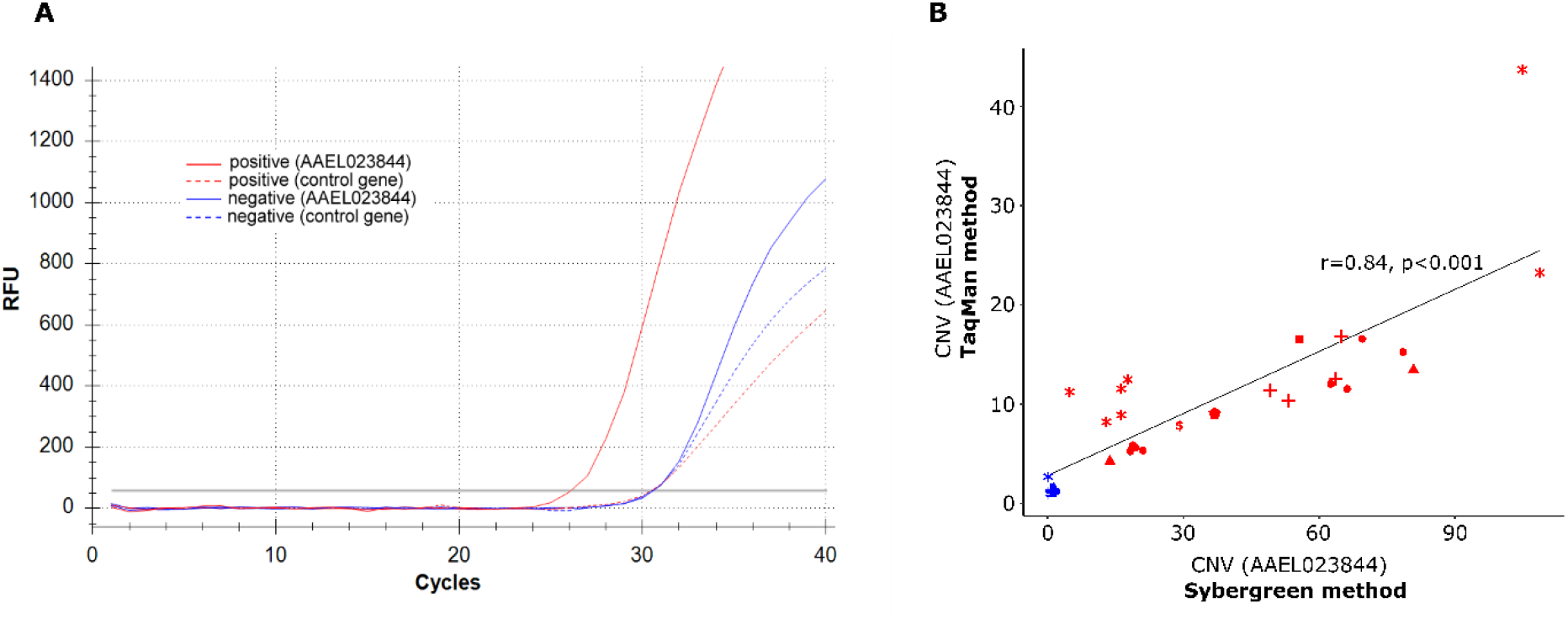
Overview of the TaqMan multiplex assay. A: Amplification profiles obtained for a positive individual (red) and a negative individual (blue). Solid line: amplification profile of the target gene AAEL023844 (formerly *CCEAE3A*, FAM probe), Dashed line: amplification profile of the control gene (AAEL007808, HEX probe). B: Comparison of CNV obtained with standard qPCR assay (SybrGreen, dual reactions) and TaqMan assay (FAM/Hex probes, single multiplex reaction). For both methods, CNV were estimated using the ΔΔCt method and are expressed as normalized gDNA quantity relative to the fully susceptible line Bora-Bora. Blue: negative individuals, Red: positive individuals. Each dot type stands for a different population.

Comparing gene copy numbers estimated from standard qPCR and TaqMan assays revealed a good correlation between the two techniques (r=0.84, P<0.001) (Figure 7B) although CNV levels obtained with the TaqMan assay were lower as compared to those obtained with qPCR and dd qPCR using amplification primers targeting a different fragment.

## Discussion

Chemical insecticides remain a key component of integrated strategies aiming to prevent the transmission of mosquito-borne diseases worldwide but the selection and spread of resistance threatens their efficacy (Moyes et al., 2017). Controlling resistance by alternating selection pressures is theoretically possible but requires an efficient monitoring of the dynamics of resistance alleles in the field (Dusfour et al., 2019). Resistance to organophosphate insecticides is common in the mosquito *Ae. aegypti* and particularly frequent in SEA following their massive use for decades (Boyer et al., 2018; Marcombe et al., 2019; Pethuan et al., 2007; Ranson et al., 2008; Vontas et al., 2012). Although several *CCE* genes are known to be involved, the genomic changes underlying resistance are not fully understood. As a result, no rapid diagnostic assay is available to monitor the frequency of resistance alleles in the field. Our experimental evolution and sequencing approaches fully supported the key role of *CCE* amplification in resistance to organophosphate insecticides in *Ae. aegypti* and allowed to further characterize the associated genomic event. The prevalence of individuals carrying this amplification was then investigated in SEA and a TaqMan multiplex qPCR assay allowing its rapid detection in single mosquito specimens was developed.

### *CCE* amplifications play a key role in organophosphate resistance in *Ae. Aegypti*

Our experimental selection approach confirmed the presence of organophosphate resistance alleles in *Ae. aegypti* populations in Laos and their rapid selection with malathion. These results are consistent with the continuous use of organophosphates for vector control for 30 years in Laos and the detection of resistance throughout the country (Marcombe et al., 2019). Bioassays with other insecticides revealed that resistance alleles selected by malathion also confer cross-resistance to other organophosphates at both larval and adult stage but not to the pyrethroid deltamethrin suggesting a resistance spectrum restricted to the organophosphate family. This confirms previous findings suggesting that the over-production of non-specific carboxylesterases is a common adaptive response to organophosphates in mosquitoes (Cuany et al., 1993; Hemingway et al., 2004; Naqqash et al., 2016). RNA-seq analysis identified seven detoxification genes over-transcribed in association with malathion resistance including five consecutive *CCE* genes on chromosome 2, one microsomal *GST* on chromosome 1, and one P450 (*CYP6N17*) on chromosome 3. Though their role in insecticide resistance cannot be excluded, the over-transcription of this *P450* and this *GST* in two other sister lines selected with insecticides from different families and showing no increased resistance to malathion does not support their key role in resistance to this insecticide (data not shown). Conversely, the over-transcription of *CCE* genes was expected as *CCE*s have often been associated with organophosphate resistance in mosquitoes. In *Cx. pipiens* their overproduction in response to organophosphate selection is well documented with distinct loci having spread worldwide (Raymond et al., 1998). In this species, high resistance levels were associated to the co-occurrence of carboxylesterases over-production through genomic amplification and the presence of the *ace-1* G119S target-site mutation affecting the acetylcholinesterase (Raymond et al., 2001). In *Aedes* mosquitoes, the G119S *ace1* mutation is submitted to a strong genetic constraint (Weill et al., 2004) and has thus not been reported, suggesting the central role of carboxylesterases over-production in resistance.

Whole genome sequencing and quantification of gene copy number supported the role of genomic amplifications in the over-production of carboxylesterases associated with organophosphate resistance in *Ae. aegypti*, as previously suggested (Faucon et al., 2015, 2017; Poupardin et al., 2014). In addition, our data revealed the co-existence of at least two distinct structural duplication haplotypes, one including the three *CCE* genes AAEL019678, AAEL023844 (formerly *CCEAE3A*) and AAEL005113 (haplotype A) and the other one not including the *CCE* gene AAEL005113 located at 5’ side of the cluster (haplotype B). Structural polymorphism of genomic amplifications in clustered detoxification genes has been recently reported in *Anopheles gambiae,* with twelve different alleles identified in a cluster of *P450*s and eleven in a cluster of *GST*s (Lucas et al., 2019). In the tiger mosquito *Ae*. *albopictus,* a structural polymorphism affecting a similar *CCE* cluster amplification was identified with at least two distinct haplotypes: one including two *CCE* genes and the second one with only the gene AALF007796, the best orthologue of *Ae. aegypti* AAEL023844 (Grigoraki et al., 2017). These striking similarities between *Ae. albopictus* and *Ae. aegypti*, likely resulting from a convergent adaptation, further supports the key role of *CCE* amplifications in the adaptation of *Aedes* mosquitoes to organophosphate insecticides.

The genetic mechanism underlying the amplification of these orthologous loci has not been characterized yet. Previous studies suggested the existence of “hot spots” of recombination favoring structural polymorphisms (Bass & Field, 2011). In insects, the presence of transposable elements is also known to favor duplication events associated with their rapid adaptation to insecticides (Bass & Field, 2011; Grigoraki et al., 2017; Schmidt et al., 2010). Our genomic data confirm the presence of multiple repeated elements in the vicinity of this locus though further genomic analyses are required to decipher their relative involvement in this genomic event.

### Evolutionary dynamics of *CCE* amplifications

The screening of this *CCE* gene amplification by qPCR on field-collected mosquitoes confirmed its occurrence in *Ae. aegypti* populations from SEA. Its prevalence in natural populations was globally low except in Cambodia and in one Thai population from which high organophosphate resistance was previously described (Paeporn et al., 2013; Pethuan et al., 2007; Poupardin et al., 2014; Saelim et al., 2005). Although our sampling campaign was restricted to a few populations in Thailand, Laos and Cambodia, the frequent elevated esterase activities detected in association with temephos resistance in SEA suggests that this *CCE* amplification is widely spread in the region (Paeporn et al., 2013; Pethuan et al., 2007). Previous studies also support the occurrence of this *CCE* amplification in the Caribbean region with high expression levels detected for AAEL023844 from multiple islands and the presence of gene amplification validated in Guadeloupe and Saint-Martin (Goindin et al., 2017; Marcombe et al., 2009, 2012). Although this needs to be confirmed, the frequent association between elevated esterase activities and organophosphate resistance in South-America (i.e. French Guiana, Brazil, Colombia and Costa-Rica) (Bisset et al., 2013; Gambarra et al., 2013; Melo-Santos et al., 2010; Paiva et al., 2016) and New Caledonia (Dusfour et al., 2015), suggests that this *CCE* amplification is distributed worldwide.

Despite the low frequency of *CCE* amplifications in most field populations, our experimental insecticide selection showed that the frequency of these resistance alleles increases rapidly in populations submitted to insecticide selection pressure. Although the potential role of genetic drift in the increased frequency of these resistance alleles in the selected Mala line cannot be fully excluded, their presence in organophosphate-resistant field populations makes it unlikely. Overall, these findings support the highly beneficial effect of these *CCE* amplifications in the presence of insecticides but also raises the question of their fitness costs in the absence of selective pressure. Fitness costs associated with the over-production of detoxification enzymes have been previously described in various insect species (ffrench-Constant & Bass, 2017; Kliot & Ghanim, 2012). Direct measurement of energetic resources (e.g. lipids, glycogen and glucose) in *Cx. pipiens* mosquitoes over-expressing carboxylesterases suggested that resistant individuals carry up to 30% less energetic reserves than their susceptible counterparts (Rivero et al., 2011). Such high metabolic cost may explain the favored selection of the shorter (less costly) haplotype B in field populations and laboratory lines showing a high *CCE* amplification prevalence. Although further studies are required to quantify the relative importance of the different *CCE* genes included in this genomic amplification in insecticide resistance, the frequent over-expression of AAEL023844 in resistant populations, its inclusion in both structural duplication haplotypes and its ability to sequester and metabolize temephos (Grigoraki et al., 2016) support its central role in organophosphate resistance.

In addition to structural polymorphism, our study revealed extensive copy number variations between resistant individuals in both field populations and laboratory lines with *CCE* gene copies varying from 3 to ~80 as measured by our TaqMan assay. In addition, no significant increase in the frequency of individuals carrying high gene copy number was observed after insecticide selection, supporting the existence of a trade-off between insecticide survival and metabolic costs associated with the over-production of these enzymes. Such high copy number polymorphism also supports the occurrence of a single duplication event followed by multiple amplification events in *Ae. aegypti* as suggested in *Cx. pipiens* and *Ae*. *albopictus* (Grigoraki et al., 2017; Guillemaud et al., 1999; Qiao & Raymond, 1995).

Though our data supported the role of this CCE amplification in organophosphate resistance, the involvement of others mechanisms cannot be excluded. Allelic variations of carboxylesterases have also been associated with organophosphate resistance in mosquitoes (Callaghan et al., 1998; Mouchès et al., 1987). Indeed, both allelic variations and genes amplifications coexist in natural populations and can be captured by selection depending on fitness-to-environment relationships (Milesi et al., 2016). In *Ae. aegypti*, a few non-synonymous variations affecting the AAEL023844 gene were associated with temephos resistance in a Thai population (Poupardin et al., 2014). However, subsequent functional studies did not support the role of these variations in insecticide sequestration and metabolism (Grigoraki et al., 2016). More recently, we combined controlled crosses with pool-sequencing to segregate organophosphate resistance alleles in a multi-resistant population from French Guiana (Cattel et al., 2019). Such approach identified a strong selection signature associated with organophosphate resistance at this *CCE* locus. Interestingly, several non-synonymous variations affecting *CCE* genes were positively associated with insecticide survival while no CNV were detected, suggesting that the selection of particular variants at this locus may also contribute to resistance. Further work is required to clarify the interplay between *CCE* amplifications, sequence polymorphism and their respective roles in insecticide resistance in *Ae. aegypti*.

### A novel TaqMan assay to track organophosphate resistance in *Ae. aegypti*

The present study supported the importance of *CCE* amplifications in insecticide resistance in *Ae. aegypti*, confirming the usefulness of CNVs for tracking resistance alleles in the field. Our cross-resistance data together with previous findings (Faucon et al., 2015, 2017; Grigoraki et al., 2016; Marcombe et al., 2019; Poupardin et al., 2014) support the routine use of this *CCE* gene amplification marker for the monitoring of resistance alleles to various organophosphate insecticides. In terms of applicability, such a CNV marker is highly superior to RNA markers classically used to detect *CCE* genes overexpression because i) genomic DNA can be extracted from dead specimens of any life stage stored at room temperature, ii) either pools or single individuals can be used if allele frequency data are required and iii) CNV quantification by qPCR is fast, easy, affordable and data are not affected by insect physiological state.

Genomic amplifications can be detected by PCR in two different ways as illustrated in Weetman et al., 2018. The first one consists of amplifying the junction between two copies by designing specific primers toward both sides of the duplicated region. Such presence/absence assay is cheap and low tech but i) does not quantify copy number, ii) requires the precise identification of duplication breakpoints, which can be impaired by the high density of repeated elements in flanking regions, and iii) may generate false negatives if duplication breakpoints vary in position or sequence. The alternative approach adopted herein consists of comparing the copy numbers between a target gene and a control gene only present as a single copy. Although this approach is slightly more expensive and requires the use of a qPCR machine, time-to-result is even shorter (no gel migration required) and results are not affected by structural polymorphisms provided an appropriate target is defined. Though multiple *CCE* genes are located within the amplified region, we selected AAEL023844 as target gene because of its central position in the genomic amplification, its inclusion in both structural haplotypes, its over-expression in several resistant populations worldwide (Dusfour et al., 2015; Goindin et al., 2017; Marcombe et al., 2019; Moyes et al., 2017) and its ability to sequester and metabolize temephos (Grigoraki et al., 2016). By targeting a coding region showing no homology with other genomic regions, we ensured a good assay specificity while limiting detrimental effects potentially caused by polymorphisms variations. Though this approach was successfully used for CNV detection with standard SybrGreen qPCR, it still required performing two distinct qPCR amplifications. Time-to-results and specificity were then further improved by the development of a dual-color TaqMan assay allowing the concomitant quantification of both target and control genes. This assay still proved to be highly specific and allowed reducing time-to-results to ~2h for less than 1.5€ /sample including gDNA extraction, qPCR consumables/reagents, amplification primers and Taqman probes.

## Conclusion

While an increasing number of alternatives to chemical insecticides are being developed for mosquito control (Achee et al., 2019) their optimization and deployment at a worldwide scale will take at least a decade. Until then, preserving the efficacy of the few insecticides authorized in public health by managing resistance is crucial to limit the impact of vector-borne diseases. However, resistance management is often hampered by insufficient resistance monitoring capacities, often leading to late or inappropriate implementations of management actions. In this context, the deep comprehension of the genetic bases of resistance and the development of molecular tools to track resistance alleles in the field still represent a significant capital gain for public health. By combining experimental selection and deep sequencing, the present study supported the key role of a genomic amplification of a carboxylesterase gene cluster in organophosphate resistance in the mosquito *Ae. aegypti*. The spatial dynamics of this resistance locus was investigated in SEA and a novel TaqMan assay was developed enabling its high-throughput monitoring in field mosquito populations. The routine use of this assay in SEA, and possibly in other tropical areas, should improve the monitoring of organophosphate resistance alleles in the arbovirus vector *Ae. aegypti*. From an evolutionary perspective, deciphering the evolutionary history of the genetic events underlying this recent adaptation undoubtedly deserves further attention.

## Data accessibility & Supplementary material

The sequence data from this study have been deposited to the European Nucleotide Archive (ENA; http://www.ebi.ac.uk/ena) under the accession numbers PRJEB37991 (RNA-seq data) and PRJEB37993 (whole genome pool-seq data). Data, scripts and supplementary material have been deposited to Zenodo (https://zenodo.org/record/4243761#.X6JBUWgzZPZ).

## Acknowledgements

The views expressed in this publication are those of the authors and do not necessarily reflect the official policy or position of the Department of the Navy, Department of Defense, nor the U.S. Government. This work was conducted in the framework of the U.S. Naval Medical Research Unit TWO projects BIO-LAO-2 (work unit number D1425) and ARBOVEC-PLUS (work unit number D1428), in support and funded by the Department of Defense Global Emerging Infections Surveillance Program and Military Infectious Disease Research Program. I (IWS and JCH) am a military Service member. This work was prepared as part of my official duties. Title 17, U.S.C., §105 provides that copyright protection under this title is not available for any work of the U.S. Government. Title 17, U.S.C., §101 defines a U.S. Government work as a work prepared by a military Service member or employee of the U.S. Government as part of that person’s official duties. This publication was also supported by the project, Research Infrastructures for the control of vector-borne diseases (Infravec2), which has received funding from the European Union’s Horizon 2020 research and innovation programme under grant agreement No 731060. Dr. Julien Cattel was supported by funding from the European Union’s Horizon 2020 Research and Innovation Programme under ZIKAlliance Grant Agreement no. 734548. The funders had no role in study design, data collection and analysis, decision to publish, or preparation of the manuscript. The field study in Cambodia was supported by ECOMORE 2 project, coordinated by Institut Pasteur and financially supported by AFD (Agence Française pour le Développement). We thank Khaithong Lakeomany, Nothasin Phommavan, Somsanith Chonephetsarath, Somphat Nilaxay, and Phoutmany Thammavong for mosquito collections and rearing from Laos. Finally, we thank Dr. Pablo Tortosa for a critical reading of this manuscript.

Version 4 of this preprint has been peer-reviewed and recommended by Peer Community In Evolutionary Biology (https://doi.org/10.24072/pci.evolbiol.100114).

## Ethical approval

Blood feeding of adult mosquitoes was performed on mice. Mice were maintained in the animal house of the federative structure Environmental and Systems Biology (BEeSy) of Grenoble-Alpes University agreed by the French Ministry of animal welfare (agreement n° B 38 421 10 001) and used in accordance to European Union laws (directive 2010/63/UE). The use of animals for this study was approved by the ethic committee ComEth Grenoble-C2EA-12 mandated by the French Ministry of higher Education and Research (MENESR).

